# Cytomixis and aberrant phenomena during meiosis in pollen mother cells of *Camellia sinensis* var. *sinensis* cv ‘Fudingdabai’

**DOI:** 10.1101/2023.04.15.537005

**Authors:** Xuelian Yu, Xinhan You, Lixia Zhang, Xingfeng Li

**Affiliations:** College of Horticulture Science and Engineering, Shandong Agricultural University, Tai’an, 271018, China; College of Agronomy, Shandong Agricultural University, Tai’an, 271018, China

**Keywords:** *Camellia sinensis*, Cytomixis, Meiosis, Microsporogenesis

## Abstract

Tea plant (*Camellia sinensis*) is an economically essential crop in China, Japan and other countries. The present study reports the meiotic behavior, including microsporogenesis of the ‘Fudingdabai’ cultivar in *Camellia sinensis* var. *sinensis*. Most of the investigated pollen mother cells undergo normal meiosis processes. In contrast, a few of the pollen mother cells showed some abnormal phenomena such as cytomixis, monovalent, laggard chromosomes, unsynchronized division, micronucleus, and so on. Among which, spontaneous cytomixis is the most common phenomenon, which mainly occurred in early prophase I but also in meiosis II. Other abnormal phenomena were less than cytomixis. The results of this study laid a foundation for exploring the meiosis and cytogenetics study of the tea plants.

## Introduction

Tea tree [*Camellia sinensis* (L.) O. Kuntze] (2n = 2x = 30) is a member of the angiosperm *Camellia* genus of the *Theaceae*, whose leaves can be used to produce different varieties of tea according to different processing processes. As a cash crop originating in the southwest of China, tea trees have played an essential role in the development of the regional economy^[1,2]^ *Camellia sinensis* var. *sinensis* cv. ‘Fudingdabai’ (FD) (accession No. GS13001-1985; Agricultural Plant Variety Name Retrieval System, Ministry of Agriculture and Rural Affairs, China), originally planted in southern China, is one of the earliest clonal tea plant cultivars recognized at the national level in China and is also the largest tea plant cultivar in China and worldwide^[3,4]^. The variety was introduced to Shandong Province in the 1980s, with strong resistance to adversity, stable quality and a wide range of adaptability, and is often used as a control variable in comparison with other varieties, so it is more meaningful to study the chromosomal behavior of pollen mother cells of ‘Fudingdabai’.

Meiosis is important in maintaining genetic stability and enriching plant variability, the abnormal phenomena during meiosis have attracted more and more attention at home and abroad. The development of cytological studies in *Theaceae* is backward, mostly focusing on analyzing chromosome ploidy and karyotype. Research on meiosis has also been carried out in several *Camellia* plants^[5]^, multiple chromosomal abnormal behaviors were found during meiosis in PMCs, such as *Camellia crassicolumna* and *Camellia oleifera* ^[6-8]^. However, for *Camellia sinensis* var. *sinensis*, the chromosome behavior during meiosis requires to be elucidated.

In this study, the flower buds of ‘Fudingdabai’ cultivar planted in Shandong Province were collected, and the microsporogenesis and pollen development process of pollen mother cells were systematically observed by pressing tablets. The purpose of this study was to provide cytological data for the pollen fertility of tea trees and to provide a reference for further studies on tea tree cytogenetics.

## Materials and methods

### Sampling and preservation

The materials are the unopened buds of ‘Fudingdabai’ planted in the “Chaxigu Planting Base” in Tai’an City, Shandong Province (116.94 E, 36.22 N). The buds measuring between 4.2 and 5.0 mm were sampled randomly from different shoots. The Buds larger than 5 mm in diameter were not used as they had passed the division stage, and 4.2 to 5.0 mm in diameter were the best source of anthers for studying meiosis.

At 8:00–10:00 am, flower buds were fixed in freshly prepared Carnoy’s fixative (Ethanol: Acetic Acid = 3:1) for 24 h, after which they were washed and stored in 70% alcohol at 4 ºC.

### Slide preparation

Both fixed and freshly collected anthers were squashed and stained with Carbol fuchsin before microscope study. Photographs of freshly prepared slides were taken with the automatic camera of Nikon Eclipse Ni-U microscope.

## Results

### Meiosis of PMCs and Microsporogenesis

Meiotic study proved that moost of the investigated pollen mother cells in ‘Fudingdabai’ undergo normal meiosis processes (Table 1). The chromosome number of ‘Fudingdabai’ was 2n = 30. Almost all the examined cells had the expected 15 bivalents ((Fig. 1f).

**Table 1.** Statistics of meiosis behavior of PMCs of ‘Fudingdabai’

| stage |  | Total<br>number<br>of cells | Normal |  | Cytomixis |  | Meiotic<br>asynchronous |  | Laggard<br>chromosomes |  | Abnormal<br>micronucleus |  | Numerical<br>abnormalities |  | Univalent |  | Unequal division |  |
| --- | --- | --- | --- | --- | --- | --- | --- | --- | --- | --- | --- | --- | --- | --- | --- | --- | --- | --- |
|  |  |  | number | frequency | number | frequency | number | frequency | number | frequency | number | frequency | number | frequency | number | frequency | number | frequency |
| Meiosis I | Leptotene | 1187 | 1182 | 99.58% | 4 | 0.34% |  |  |  |  | 1 | 0.08% |  |  |  |  |  |  |
|  | Prophase | 1381 | 1324 | 95.87% | 50 | 3.62% |  |  |  |  | 6 | 0.43% |  |  |  |  | 1 | 0.07% |
|  | Pachytene | 147 | 146 | 99.32% |  |  |  |  |  |  |  |  |  |  |  |  | 1 | 0.68% |
|  | Diplotene | 176 | 167 | 94.89% | 6 | 3.41% |  |  |  |  |  |  | 3 | 1.70% |  |  |  |  |
|  | Diakinesis | 439 | 407 | 92.17% | 12 | 2.73% |  |  |  |  |  |  | 16 | 3.64% | 4 | 0.91% |  |  |
|  | Metaphase I | 657 | 613 | 93.30% | 27 | 4.11% |  |  | 8 | 1.22% |  |  |  |  | 9 | 1.37% |  |  |
|  | Anaphase I | 67 | 65 | 97.01% |  |  |  |  | 2 | 2.99% |  |  |  |  |  |  |  |  |
|  | Telophase I | 206 | 202 | 98.06% |  |  |  |  | 4 | 1.94% |  |  |  |  |  |  |  |  |
| Meiosis II | Prophase II | 538 | 478 | 88.85% | 13 | 2.42% | 14 | 2.60% | 2 | 0.37% | 9 | 1.67% |  |  |  |  | 22 | 4.09% |
|  | Metaphase II | 399 | 338 | 84.71% | 37 | 9.27% | 13 | 3.26% | 4 | 1.00% | 5 | 1.25% |  |  | 2 | 0.50% |  |  |
|  | Anaphase II | 394 | 356 | 90.36% | 14 | 3.55% | 6 | 1.52% | 5 | 1.27% | 2 | 0.51% | 9 | 2.28% | 2 | 0.51% |  |  |
|  | Tetrahedral II | 1581 | 1481 | 93.67% | 78 | 4.93% |  |  |  |  | 2 | 0.13% | 15 | 0.95% |  |  | 5 | 0.32% |
|  | Tetrad | 415 | 415 | 100% |  |  |  |  |  |  |  |  |  |  |  |  |  |  |
| Total |  | 7587 | 7174 | 94.56% | 241 | 3.18% | 33 | 0.43% | 25 | 0.33% | 25 | 0.33% | 43 | 0.57% | 17 | 0.22% | 29 | 0.38% |

**Fig. 1.**
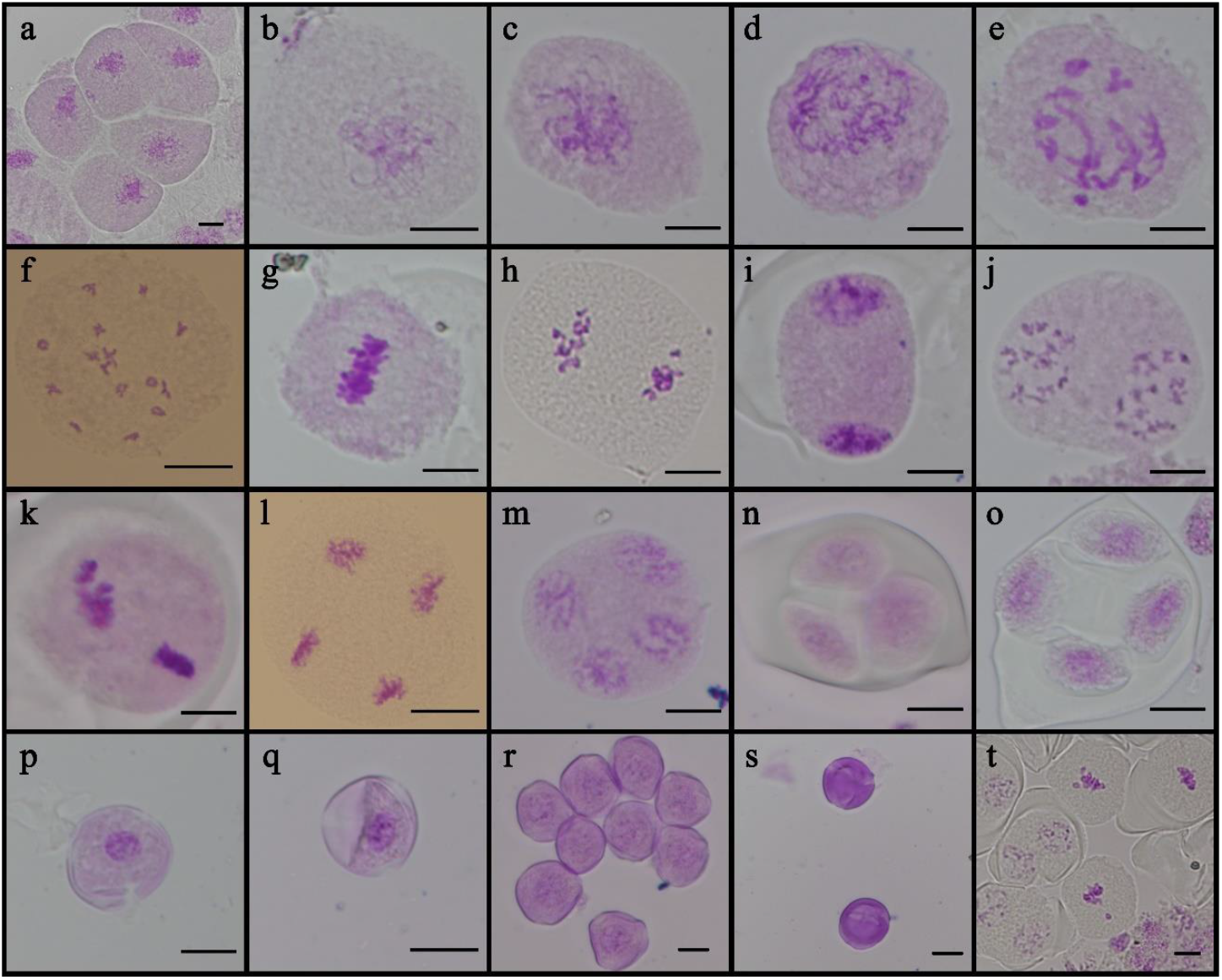
The microsporogenesis of PMCs in *C. sinensis* (a) Early leptotene, (b) Leptotene, (c) Zygotene, (d) Pachytene, (e) Diplotene, (f) Diakinesis, (g) Metaphase Ⅰ, (h) Anaphase Ⅰ, (i) Telophase Ⅰ, (j) Prophase Ⅱ. (k) Metaphase Ⅱ. (l) Anaphase Ⅱ, (m) Telophase Ⅱ, (n) Tetrahedral tetrad, (o) Planar tetrad, (p) Monokaryon, (q) Late uninucleate stage, (r) Immature pollen, (s) The mature pollen, (t) Multiple meiotic phases were observed in one flower bud. Bar=10μm

At the beginning of meiosis, PMCs with dense cytoplasm huddled together in the anther and were challenging to separate. In the early leptotene stage (Fig. 1a), filiform chromosomes were irregularly distributed around the nucleolus, where the cytoplasm was still thick. Chromosomes gradually gathered at one side of the nucleolus during the late leptotene stage (Fig. 1b), which presented as a truss. Homologous chromosomes pairing together, and partial juxtaposition of the chromosomes in the cytoplasm can be observed around the nucleolus (Fig. 1c). The chromosomes continued to coarsen during the pachytene stage (Fig. 1d), and chromosomes remained entangled. During the diplotene stage (Fig. 1e), homologous chromosomes began to repel each other, although chromosome chiasmata were observed. Chromosomes were highly coagulated, and 15 bivalents separated equally in the cell, facilitating chromosome counting during the diakinesis stages (Fig. 1f). At metaphase Ⅰ (Fig. 1g), chromosomes were neatly established along the equatorial plate. Subsequently, during anaphase Ⅰ (Fig. 1h), chromosomes of homologous pairs were pulled to different poles of the cell(Fig. 1i). During the second division, the chromosomes became diffuse at prophase Ⅱ (Fig. 1j), arranged themselves properly on the equatorial plane at metaphase Ⅱ (Fig. 1k), and chromatids moved to the opposite poles during anaphase Ⅱ (Fig. 1l) and nuclei reformed at telophase Ⅱ (Fig. 1m). Finally, the tetrads were formed. Cytokinesis was simultaneous, and the microspore tetrads were tetrahedral (Fig. 1n, o).

After meiosis, the microspores are released and developed with dense cytoplasm. The nucleus was located in the center of the cell with a large volume in mononuclear microspore stage (Fig. 1p). With the development of mononuclear microspores, vacuoles appear in the cell and push the nucleus towards the cell wall. This stage is the sidelining stage of mononuclear microspores (Fig. 1q). Then, the microspores begin to undergo mitosis and form pollen grains as the microspores mature (Fig. 1r, s). Multiple developmental stages of microsporogenesis were observed in one bud (Fig. 1 t).

### Cytomixis in ‘Fudingdabai’

Among all the abnormal phenomena in microsporogenesis, spontaneous cytomixis appeared with the higher frequently in ‘Fudingdabai’. Cytomixis involving the transfer of chromatin material via the cytoplasmic channels among adjacent PMCs at various stages of meiosis was observed in *C. sinensis* (Fig. 2), which mainly occurred at prophase Ⅰ (10.1%), followed by M Ⅱ (9.27%).

**Fig. 2.**
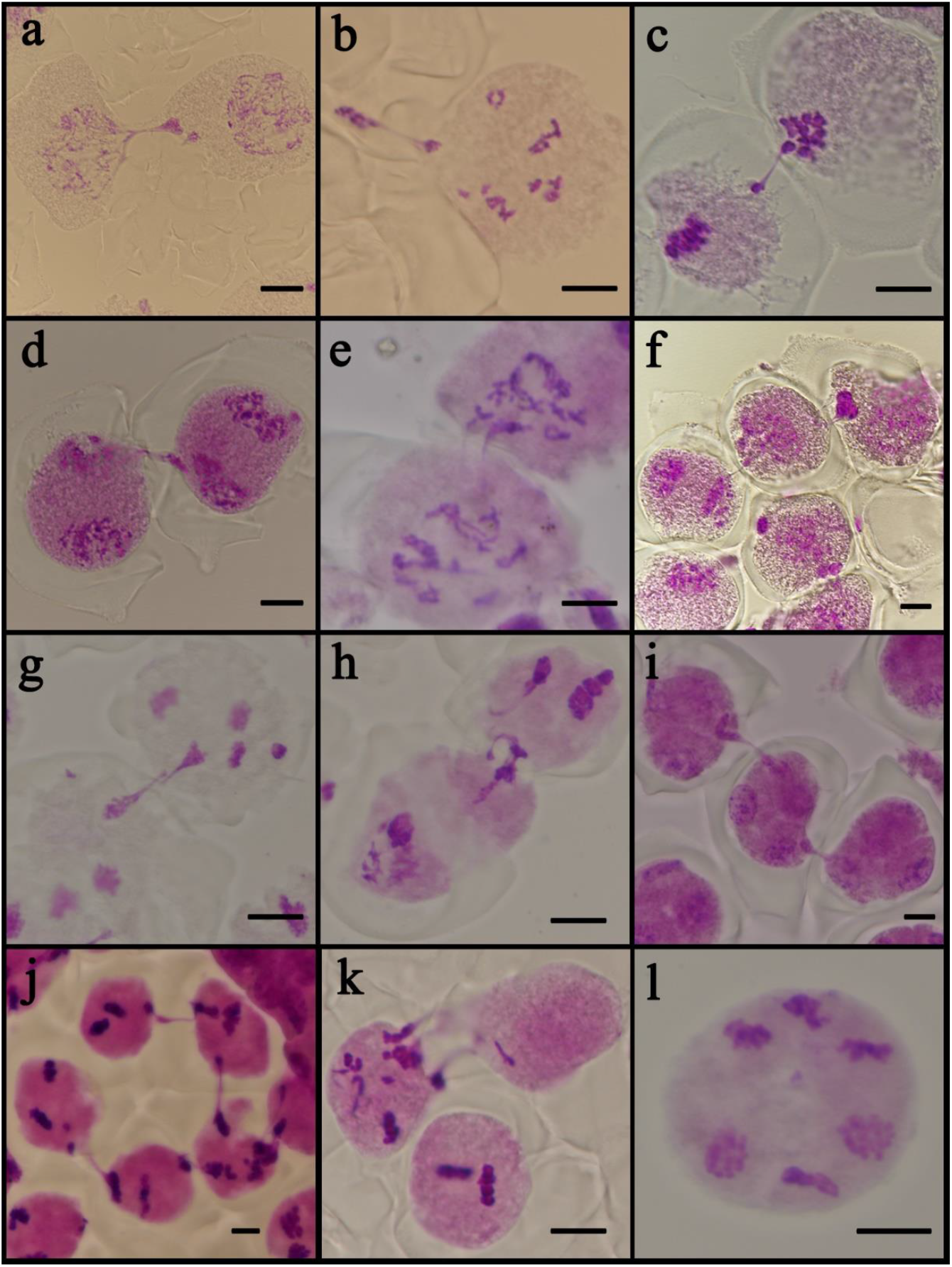
Cytomixis in ‘Fudingdabai’ (a-d) Cytomixis in PMCs at a different stage, (e) Cytomixis in PMCs through more than one cytomictic channels, (f) Chromosome migration in PMCs between different stages, (g-j) the recipient cell passes the chromatin on to another cell sometimes, (k) the whole nucleus migrates, (l) hyperploid PMCs. Bar=10μm

At the leptotene stage of prophase Ⅰ, the chromatin were condensed and visible as a chromatin balls that lay close to the cell membrane with clustered cytomictic channels (CCs) (Fig. 2a-d). Chromatin migration caused the formation of single or multiple chromatin bridges between PMCs (Fig. 2e). The chromatin might transferred either in one direction or randomly. One cells usually transferred chromatin to one of the neighboring cells, but sometimes also to two or more cells, or even from two or more nuclei into one cell. Chromatin migration not only happen between cells that were in the same stage, but also cells in different stages of meiosis (Fig. 1f). In some instances, the recipient cell passed the chromatin on to another cell sometimes, it appeared as though chromatin passed from the first meiocyte to the second, from the second to the third, and so on (Fig. 2g-j). The amount of chromatin being transferred ranged from one chromosome to some chromosomes, but it rarely happened that the whole nucleus would migrate (Fig. 2k), leaving the meiocyte de-nucleated and resulting in hypo- and hyperploid PMCs (Fig. 2l).

### Other abnormal chromosome behaviors

Apart from cytomixis, other aberrant chromosome behaviors were also observed. These induced the presence of laggard chromosomes, chromosome bridges, univalent, unequal clusters of chromosomes and fragments in microsporogenesis (Table 1).

Several chromosomes were not synchronized (neatly arranged on the cell plate) in metaphase Ⅰ (Fig. 3a). During the observation of anaphase Ⅰ in ‘Fudingdabai’ PMCs, it was observed that some cells had lagging chromosomes (Fig. 3b), which showed that when chromosomes were pulled to both poles by spindle filaments, a few chromosomes did not move to both poles or moved more slowly. Similarly, chromosome separation was not synchronized in anaphase Ⅱ (Fig. 3c), and lagging chromosomes were observed (Fig. 3d). These lagging chromosomes or fragments would formed the micronuclei laterly. The presence of micronuclei was found at prophase Ⅰ (Fig. 3e) and prophase Ⅱ (Fig. 3f). Univalent were also observed in some cells at diakinesis (Fig. 3g). Sometimes the chromosomes in different dyad would undergone different stages of division (Fig. 3h-j). Another well-known phenomenon was abnormal karyokinesis and cytokinesis in PMCs (Fig. 3k-n), resulting in unequal distribution of chromosomes/chromatin in daughter cells. All these abnormalities can lead to an increased (Fig. 3o) or decrease (Fig. 3p) of chromosomes.

**Fig. 3.**
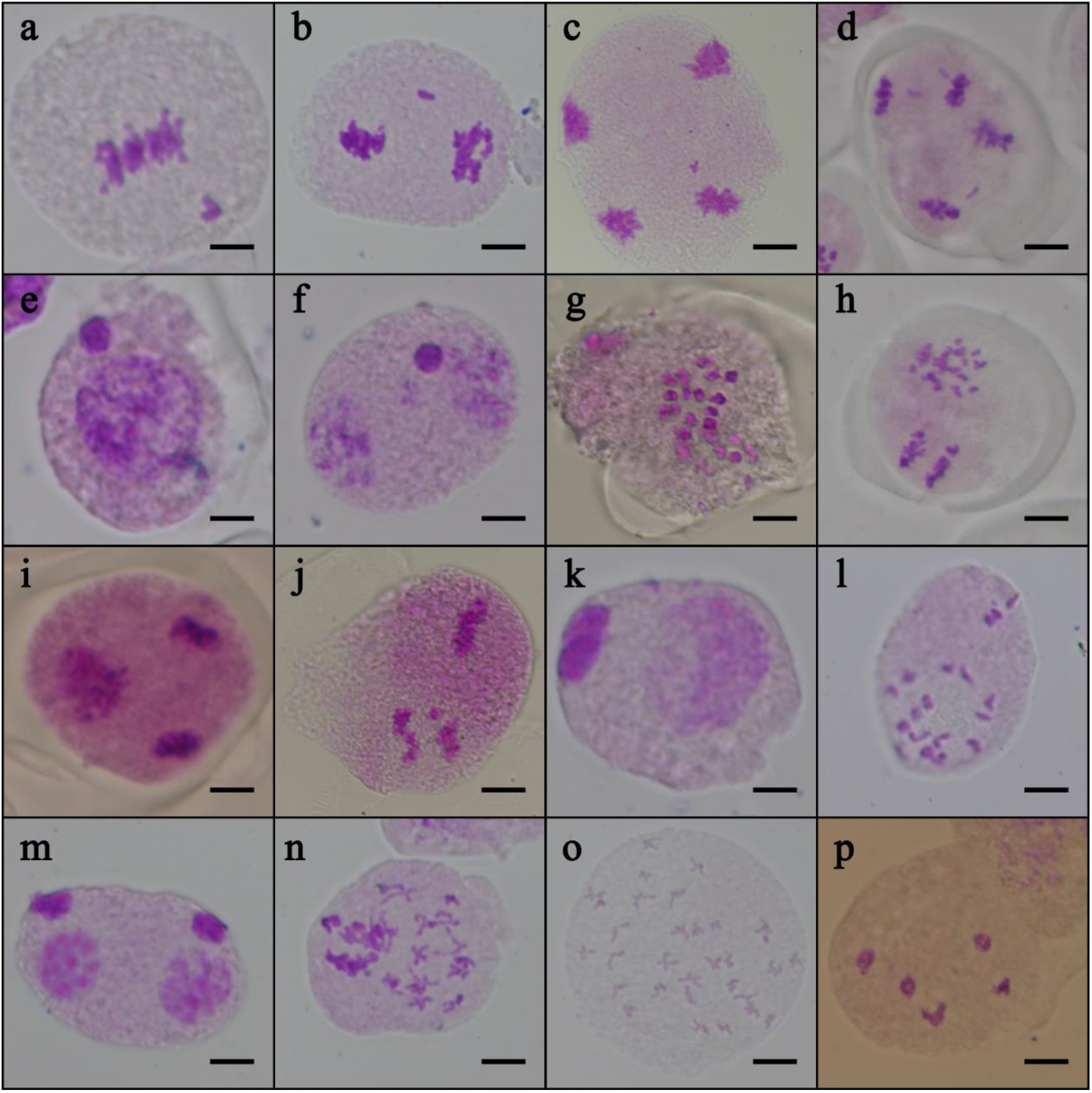
Other abnormalities in meiosis of ‘Fudingdabai’ PMCs (a) Chromosomes were not synchronized in metaphase Ⅰ, (b-d) Lagging chromosomes, (e-f) Micronuclei, (g) Univalent chromosome, (h-j) Chromosomes on either side of the cell are at two different stages of division, (k-n) Abnormal karyokinesis and cytokinesis, (o) Increase of chromosomes, (p) Decrease of chromosomes. Bar=10μm

## Discussion

The present study observed male meiosis and microsporogenesis from ‘Fudingdabai’ of *C*.*sinensis* in the Shandong Province of China. The results showed that meiotic abnormalities, including cytomixis, laggard chromosomes, micronuclei, and chromosome separation, were not synchronized and appeared in about 5.44% of PMCs, and abnormal cytokinesis (Table 1, Figures 2, 3). Abnormal meiotic processes often result in disruption of microsporogenesis, leading to pollen aberrations or sterility and negatively impacting the reproductive success of the species^[9-12]^.

Cytomixis is one phenomenon that is under the influence of genetic and environmental factors^[13]^. After it was first recorded by Arnoldi (1900), the phenomenon of cytomixis and other abnormalities responsible for abnormal meiotic behavior and reduced pollen fertility have been reported in some flowering plants^[9,12,14-16]^. However, the term ‘cytomixis’ was coined by Gates (1911), who studied the PMCs of *Oenothera gigas* and defined it as a phenomenon of transmigration of chromatin from one cell to an adjacent cell Cytomixis involves the migration of chromatin material among meiocytes through cytomictic connections^[17]^. This phenomenon is most frequently observed in male meiosis and has been so far described in the microsporogenesis of over 400 higher plant species ^[18,19]^. In many of the earlier studies, cytomixis had been held directly responsible for inducing abnormal meiotic behavior, pollen sterility, and pollen grains of variable sizes^[20,21]^.

Many authors have suggested cytomixis as an artifact of fixation^[22,23]^. To reduce the effect of fixative on cytomixis, freshly collected anthers were stained immediately for observation without fixation, yet there appeared to be no difference in the incidence and frequency of cytomixis between fresh and fixed anthers. This result dramatically excludes fixation as a factor inducing cytomixis in the present study. Many authors previously reported this finding^[21,24]^.

Cytomixis has been reported to be observed only at the early stages of the first meiotic division in many plants^[11]^, whereas in *Camellia sinensis* cell fusion was observed at every stage, even at the telophaseⅡ. In most cases, the nucleus does not pass to the recipient cell as a whole. Once the CC is passed, parts of the nucleus bud off to form one or several micronuclei in the recipient cell cytoplasm, whereas the more significant part of the migrating nucleus remains in the donor cell. When a whole nucleus migrates to the recipient cell to form, a binucleated meiocyte is rarer^[25,26]^.

Aberrant chromosome behavior other than cytomixis was also observed, but the frequency was much lower than cytomixis. The number of meiotic abnormalities, such as univalents, bridges, fragments, unequal separation, and lagging chromosomes, was recorded and analyzed at different stages. Due to cytomixis, a change in the amount of chromatin or the number of chromosomes in the cells is involved. Depending on the nature of cytomixis, cells with the aneuploid number of chromosomes are formed probably through cytoplasmic connection type. Micronucleus formation was caused by chromosome lag that prevented chromosomes from entering newly formed cells^[27]^. The formation of lagging chromosomes is a product of several pre-anaphasic chromosomal irregularities^[28]^. However, the detailed mechanisms underlying these observations in *C. sinensis* are unclear.

In conclusion, we observed a high frequency of aberrant phenomena in the meiosis of ‘Fudingdabai’, with the highest frequency of cell fusion occurring at almost every period, unlike in other plants where it was reported to occur only in the meiosisⅠ. Other anomalies occurred much less frequently than cell fusion. However, more work is needed to elucidate the true causes, consequences and significance of the occurrence of meiotic anomalies such as cell fusion in tea trees.

